# Proactive inhibition of goal-directed movements involves explicit changes to movement planning

**DOI:** 10.1101/2022.06.01.494317

**Authors:** John P. Pickavance, M. Mon-Williams, Faisal Mushtaq, J. Ryan Morehead

## Abstract

Inhibition can be implemented reactively, withholding movement in response to a stop-signal, or by proactive changes to movement planning when a stop-signal is expected. Previous studies have typically employed simple button presses, finding proactive delays to movement onset when a stop-signal might appear. Here, we consider inhibition in the context of more complex, goal-directed movements, such as the swing of a bat. Thus, we observe two additional dimensions of movement planning under proactive control, movement duration and end-point error. We found, in addition to onset delay, movements were briefer and arrived later when a stop-signal might appear. This challenges a classical theoretical dichotomy, suggesting proactive inhibition is underlay by both response suppression and delayed facilitation. Moreover, participants were aware of delays to onset and arrival, but reported magnitudes were smaller than observed. This suggests proactive inhibition operates as an explicitly retrievable compensatory strategy whose finer details are implicitly tuned.

## INTRODUCTION

In baseball, batters must not only plan and execute an accurately timed swing, but also inhibit movement in response to an errant pitch. Successful inhibition can be implemented reactively, driven by a stop-signal^1^ (e.g., the pitch veering off course), or proactively, by preparing to stop in anticipation of a stop-signal^2^. Reactive inhibition has been explored in a range of contexts, including simple reaction times^1,3^, choice reaction times^4,5^, anticipation timing^6–8^, and interceptive actions^9^. These studies have shown that people are able to inhibit prepotent movements in response to stop-signals approximately 200ms prior to their initiation. This measure of inhibition, the stop-signal reaction time (SSRT), has been critical to understanding a range of neuropathologies because it is often elevated in conditions characterized by inhibitory deficiencies, such as Parkinson’s disease^10^, attention deficit/hyperactivity disorder^11–13^, addiction^14^, and obsessive compulsive disorders^15^.

Movements can be proactively inhibited by delaying their onset, which increases one’s chance of detecting a need to stop before initiation^7,16–18^. In contrast to reactive inhibition, which is automatic, proactive inhibition is thought to be volitional and strategic^2,19^. Accordingly, it has been argued that proactive control may represent the more ecological dimension of inhibition. For example, an addict’s substance avoidance or the suppression of a child’s tics is unlikely to recruit reactive mechanisms in response to external stimuli, but may depend rather on the ability to delay their expression^2^. Nevertheless, there is some debate as to whether inhibition is implemented cognitively at all^20^. On the one hand, delays to movement onset under stop-signal uncertainty may reflect response suppression (or cognitive inhibition)^2,21,22^, on the other, it may simply reflect delayed facilitation (or a “waiting” strategy)^23^. Our understanding of this phenomenon is mostly limited to simple button press tasks in which response delay is the only signature of proactive inhibition. We believe it is possible to tease apart the underlying mechanisms of inhibition by considering inhibitory control in the planning and execution of more sophisticated behaviors, such as the swing of a bat.

To examine proactive control, we adapted a task previously used to investigate reactive inhibition in the context of interceptive actions^9^. In this paradigm, the interception point is fixed and the timing characteristics of the movement can be considered in isolation^24^. Thus, in addition to movement onset (initiation time), this paradigm allows us to investigate the duration of movement (movement time) and the associated end-point error (timing error). Previous interception studies have shown timing error remains fixed over a range of timing constraints, with any delays in initiation offset by decreases in movement time^25–27^. It is thought movement times are programmed in advance based on the required precision, and triggered when estimates of the time taken for the target to reach the interception point are equivalent (plus neurophysiological delays)^24^. According to this scheme, if the triggering of the prepared response is suppressed (i.e., takes longer to complete^16^), the prepared movement time will be inappropriate to preserve timing error and arrival to the target will be delayed. Alternatively, any delays in facilitation should be offset by a proportional reduction in movement time because it is specified immediately prior to triggering^9^, and can be programmed to reflect this wait.

By comparing the characteristics of timed interceptive movements between conditions in which stop-signals could be presented (Uncertain) and in which they were never presented (Certain), we were able to investigate the mechanisms underlying proactive delays to initiation. We found evidence for both delayed facilitation (shorter movement times) and response suppression (delayed arrival). Notably, these timing characteristics can be expressed spatially, with the position of the target at initiation specifying initiation time and the point of the target hit specifying timing error. This allowed us to quantify the degree to which participants were explicitly aware of their strategies using a spatial assay. We show participants were aware of delays to initiation and arrival, but this does not fully account for the magnitude of the effects we observed. Collectively, these results challenge classical theories of inhibition and present a new way forward for examining the processes underlying proactive control.

## RESULTS

### Task performance met the assumptions of the independent race model of inhibition

We first validated our task, considering whether sensible values of SSRT could be estimated and that behavior was suitably constrained by the presentation of stop-signals. Participants tried to hit a target as it moved overhead with a bat controlled by mouse or trackpad (Fig. 1A). When the background showed dawn (indicated by the color of the sky), there was a 33% chance a stop-signal (instantly switching to daytime; Fig. 1B) could be presented. Participants were asked to withhold their movement in response to the stop-signal (stop-trial) and hit the target in its absence (go-trial). On the subsequent stop-trial, the stop-signal was delayed (cf. brought forward) an additional 50ms if participants successfully inhibited (cf. incorrectly responded).

**Figure 1.**
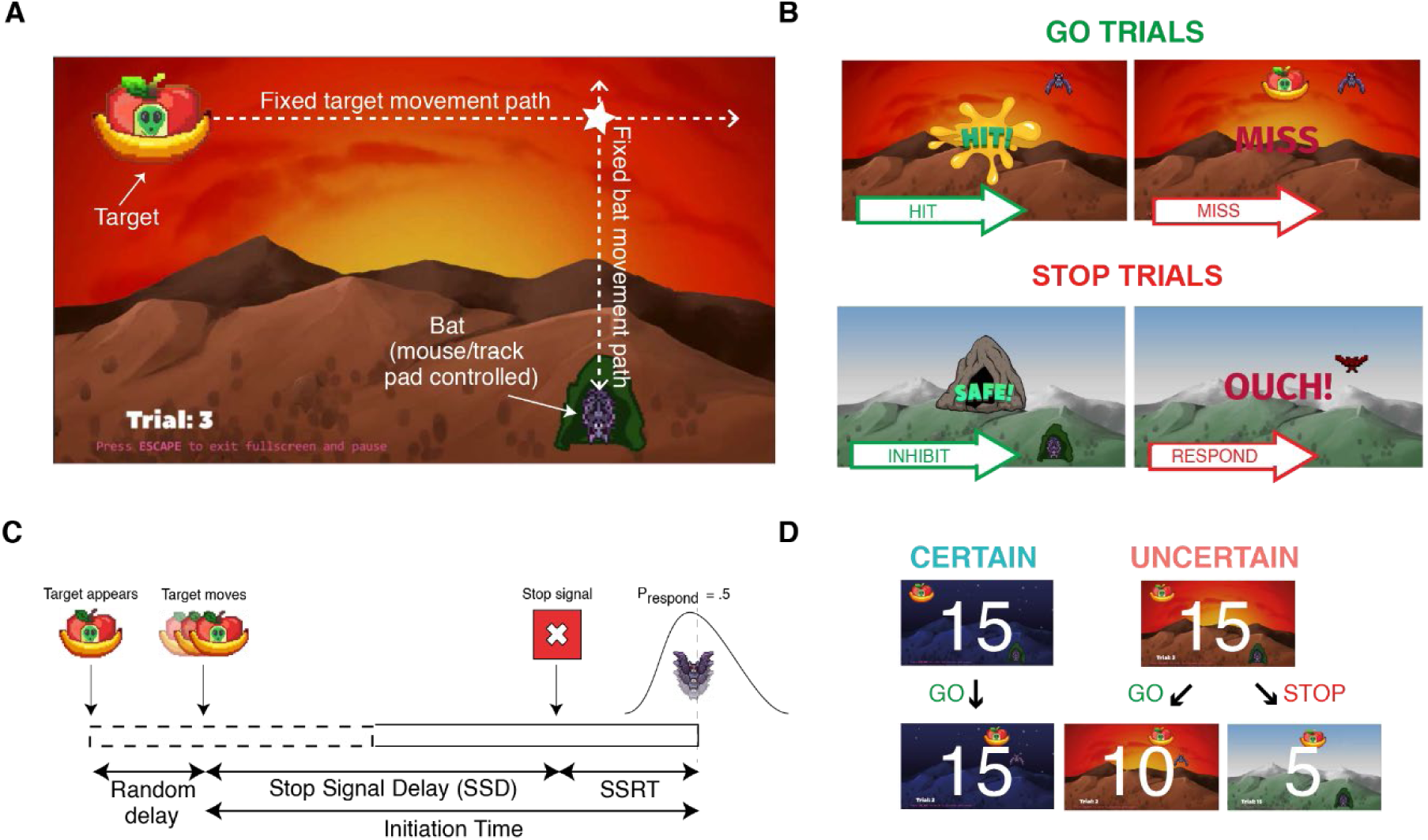
Task Schematic. (A) The target appears in the top-left of the screen. After a random delay, it moves from left to right. (B) On Go trials, participants try to hit the target with the bat. Feedback on hit success was given after the movement. On Stop trials, a stop-signal was denoted by the scene switching to daytime. Feedback was given depending on whether movement was successfully inhibited. (C) The Stop-Signal Reaction Time (SSRT) was calculated from estimates of the mean initiation time on Go trials and the SSD at which the probability of successfully inhibiting a movement was 0.5. (D) Experimental Blocks. Each block comprised 15 Certain (night) and 15 Uncertain (dawn) trials. There were 8 blocks, yielding 240 trials total. Within each block, the order of trials was random. Uncertain trials were used to estimate SSRT. The difference between initiation time, movement time, and timing error, on Certain and Uncertain Go trials, comprised our measures of proactive inhibition. At the end of the task, after all movement trial blocks, participants reported where the target was when they initiated movement, and where they aimed to hit the target, for both the Certain and Uncertain conditions.

SSRTs were calculated using the integration method. These estimates rely on assumptions made by the independent race model of inhibition to be met. The model states inhibition is a race between a go process which begins accumulating in response to the imperative stimulus (i.e., the target reaches a critical point in its trajectory^9^), and a stop process which accumulates in response to the stop-signal (i.e., the switch to daytime).

We empirically verified the applicability of the independent race model with four key observations^28^. Firstly, failures to stop should be earlier than responses on go trials because the stop process cuts off the upper tail of the go initiation time distribution (Fig. 2A). Secondly, if longer SSDs handicap the race in favor of the go process, the probability of correctly inhibiting should increase with earlier SSDs (Fig. 2B). Similarly, failed stop movement amplitudes should increase with later SSDs because movements will be interrupted later in their course (Fig. 2C). Finally, taken together, stop failures should be earlier with earlier SSDs because only the quickest go processes can outrace the earliest accumulating stop processes (Fig. 2D). Having confirmed these assumptions empirically, we can be confident in our estimates of SSRT derived from it. In our interceptive stop-signal task, the mean SSRT was 197ms (n = 50, 95%C.I. = 13.3ms).

**Figure 2.**
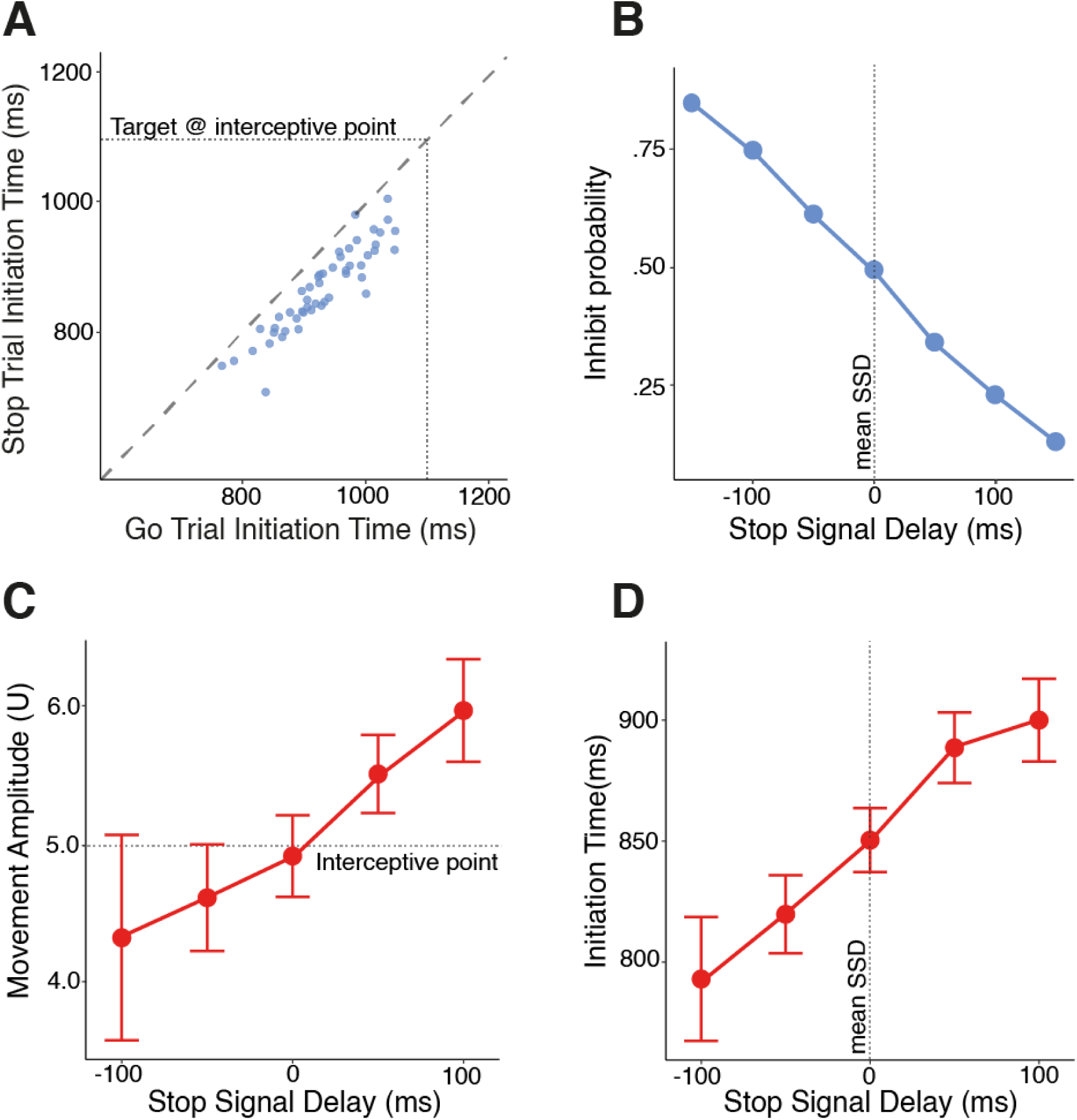
Empirical verification of the applicability of the horse race model. (A) Initiation times on go trials were longer than initiation times on failed stop trials for all participants. Each blue dot represents a single participant. The oblique, dashed line of equilibrium shows where initiation times would be equal. (B) The probability of correctly inhibiting decreases with longer stop-signal delays. Each blue dot represents the proportion of trials inhibited from the total at each level of stop-signal delay. (C) The amplitude of incorrect movements on stop trials increases with longer stop-signal delays. (D) Initiation times on failed stop trials increased with longer stop-signal delays. To allow direct comparison, stop-signal delays have been centered for each participant by subtracting their mean SSD. For panels (C) and (D) each red dot represents the population mean at each level of stop-signal delay, with error bars representing 95% confidence intervals.

These findings, in addition to the summary statistics presented in Table 1^28^, give us confidence in the validity of our SSRT estimates. Nevertheless, delays to initiation can bias estimates of SSRT. Considering the average of binned estimates of SSRT in two blocks^29^, our estimate of the mean SSRT was 197ms(± 13.7) and not significantly different from the original estimate (*t*(49) = -.581, *p* = .564). Additionally, mean go-trial responses times can be quicker than incorrect responses on stop-trials they immediately precede, particularly at the shortest SSDs^30^. In our task, however, mean initiation times on stop-trials were quicker than on go-trials at all SSDs (Supplement C). In summary, the independent race model of inhibition holds. We can be confident participants performed the task as intended, thus movements on go-trials could be suitably constrained by the uncertainty of whether a stop-signal would be presented^16^.

**Table 1.** Summary statistics of measures used to calculate stop-signal reaction times

|  | Mean | SD | n |
| --- | --- | --- | --- |
| Probability of no response on go-trials | .021 | .021 | 50 |
| Probability of hitting target on go-trials | .924 | .064 | 50 |
| Initiation Time on go-trials (ms) | 931 | 69.4 | 50 |
| Probability of inhibiting on stop-trial | .500 | .037 | 50 |
| Nth trial | 39.5 | 2.96 | 50 |
| Mean stop-signal delay (ms) | 738 | 88.5 | 50 |
| Stop-signal Reaction Time | 197 | 46.9 | 50 |

### Stop-signal uncertainty led to briefer movements with delays to initiation and arrival

Concurrently, we assessed the effects of stop-signal uncertainty on interceptive movements. In our task, participants were told it was always safe to leave the cave at night (Certain), whereas the sun may come out at dawn (Uncertain) and movement should be withheld if it does. Each block consisted of 15 Certain (15 go-) and 15 Uncertain (10 go-, 5 stop-) trials and there were 8 blocks in total for 240 trials (Fig. 1D). Trial order was randomized within block and all participants experienced the same order. Thus, we compared movement characteristics between Certain and Uncertain go-trials to ascertain the effects of proactive inhibition.

Our task yielded three measures of proactive inhibition: initiation time; movement time; and timing error. Initiation times were converted to time to contact (TTC) values by subtracting the observed initiation time from the time taken for the target centre to reach the interceptive point (i.e., 1100ms). Larger values of TTC therefore represent initiations that are earlier in the target’s trajectory, and any discrepancy between TTC initiation time and movement time will result in a timing error of equivalent size. All three measures of proactive inhibition demonstrated non-linear changes over the course of the experiment (Fig. 3A). Accordingly, 3-parameter exponential models were fit for each measure that allowed the intercept and asymptote to vary by Certainty (with random slopes and intercepts on both parameters). Contrasts were performed on mean estimates produced by the models by Certainty for each participant, with the intercept approximating performance at the start of the experiment and the asymptote approximating performance at the end. Thus, we were able to quantify proactive inhibition by considering differences in initiation time, movement time, and timing error between Certainty conditions, at the start and end of the experiment.

**Figure 3.**
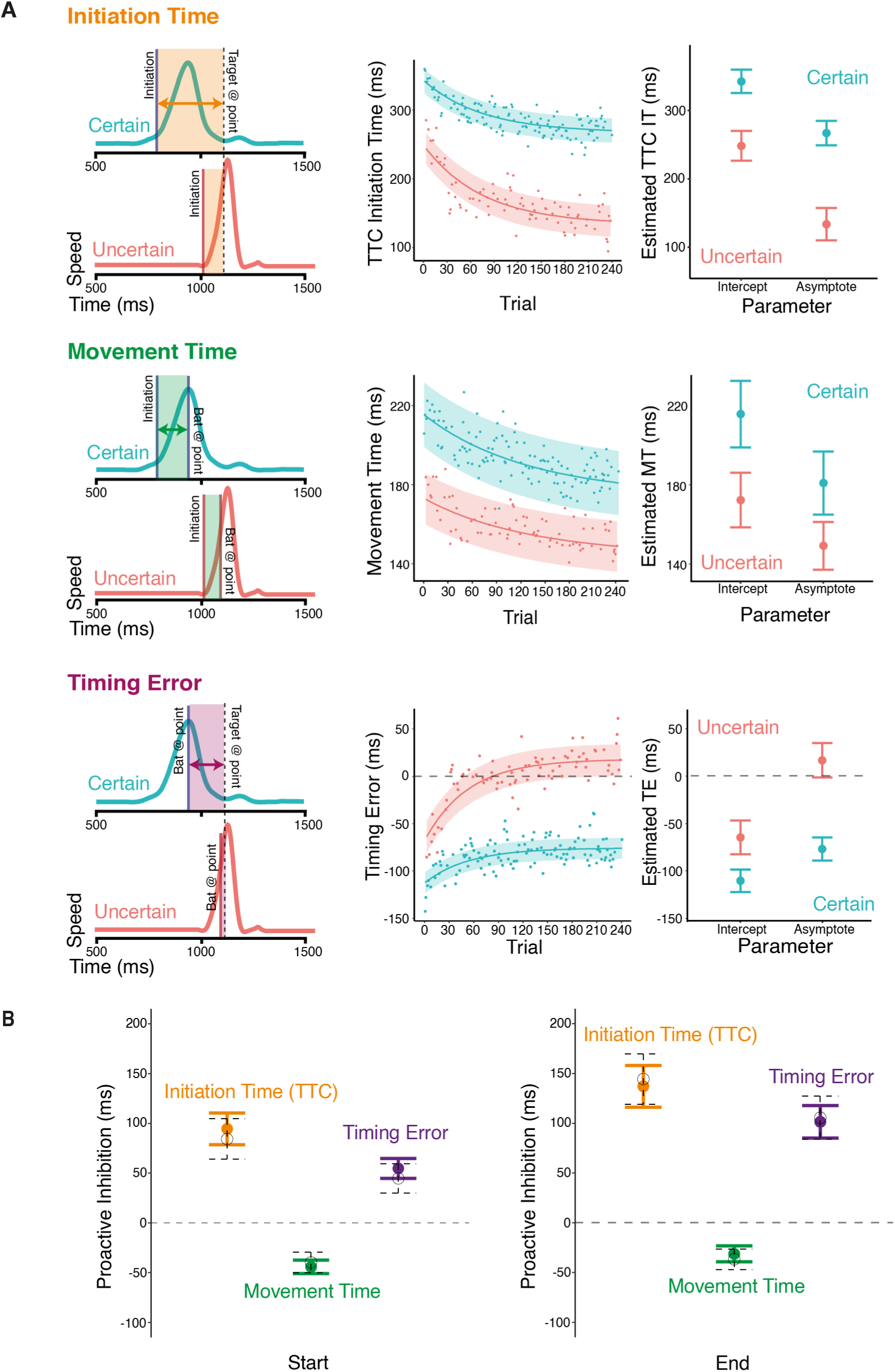
A) Quantifying the effects of stop-signal uncertainty on interceptive movements by comparing kinematic measures between Certain (blue) and Uncertain (red) conditions. Top: (Left)Velocity-time profiles of representative movements in each condition. The initiation line indicates when the movement was initiated (i.e., displaced 0.7U), the bat at point line represents when the bat reaches the interceptive point (i.e., displaced 5U), and the target at point line represents when the center of the target would reach the interceptive point (i.e., after 1100ms). The colored/shaded areas represent the time elapsed for each measure. TTC Initiation Time (orange) is the time elapsed from initiation until the center of the target would reach the interceptive point. Movement Time (green) is the time elapsed from initiation until the bat crosses the interceptive point. Timing Error (purple) is the difference in time between when the bat reaches the interceptive point and the center of the target would reach the interceptive point (negative is earlier, positive is later). (Middle) 3-Parameter asymptote model fits for each measure. Individual points represent the observed population mean for each trial. Solid lines are estimates of the mean, with the shaded area representing 95% confidence intervals. (Right) Mean estimates of intercept and asymptote parameters by certainty. Points represent population mean, with error bars indicating 95% confidence interval. B) Each measure of proactive inhibition represents the difference in timing between Certain and Uncertain conditions. Coloured dots (error bars) represent the mean (95% confidence interval) of estimates extracted from our model parameters for each participant. Participants delay their initiation and quicken their movement in response to stop-signal uncertainty. Movements are not sufficiently quicker to preserve timing error, thus arrival to the target is also delayed. Delays to initiation and arrival are more pronounced at the end of the experiment. Black dots (dotted lines) show the mean (95% confidence interval) of participants’ model-free proactive differences which are not statistically different from our model estimates.

Initiation was delayed in response to stop-signal uncertainty (F(1, 49) = 202.56, p < .001), and as the experiment progressed (F(1, 49) = 81.54, p < .001). The interaction (F(1, 49) = 18.58, p < .001) suggests participants delayed more in the Uncertain condition as they went on (Fig. 3). Considering the differences, participants initiated 95(±16)ms later in Uncertain relative to Certain conditions at the beginning of the experiment, and 134(±21)ms later by the end (Fig. 3A).

Movement time was similarly affected by uncertainty (F(1, 49) = 159.39, p < .001) and progress (F(1, 49) = 25.24, p < .001). There was also an interaction (F(1, 49) = 4.78, p = .034) indicating participants shortened movement times less in Uncertain conditions over the course of the experiment (Fig. 3). In terms of the overall difference, participants moved 44(±7)ms quicker in Uncertain relative to Certain conditions, which diminished to 33(±8)ms by experiment end (Fig. 3A).

The shortening of movement time was not sufficient to preserve timing error, given the delays to initiation. Indeed, there were effects of Certainty (F(1, 49) = 193.36, p < .001) and progress (F(1, 49) = 81.11, p < .001) on timing error, and an interaction between the two (F(1, 49) = 32.88, p < .001) such that participants delayed arrival more in Uncertain conditions as the experiment progressed (Fig. 3A). Proactive differences derived from the mean estimates show participants initially arrived 54(±10)ms later to the target in the Uncertain condition, which increased to 99(±16)ms over the course of the experiment (Fig. 3).

To be sure, we additionally considered a model-free analysis comparing participant means of each measure from the first and last experimental block. Mean and model estimates of proactive differences in initiation time, movement time, and timing error, at both the start and end of the experiment, were indistinguishable (Fig. 3B). In summary, participants delayed initiation and shortened movement time in response to stop-signal uncertainty, but the movement time shortening was not sufficient to preserve timing error and participants arrived later to the target. These differences were greater at the end of the experiment, with later initiation accompanied by less shortening of movement time, producing later arrival to the target in Uncertain conditions.

### Participants were explicitly aware of proactive changes to their movements

We found participants delayed their movement initiation and their arrival to the target in response to stop-signal uncertainty. But does this represent a volitional strategy? In motor control studies self-reported measures of aim are commonly used to quantify the extent changes in behavior are governed by explicit processes^31,32^. In our task, the interceptive point is fixed, and the target moves at a constant speed. Accordingly, initiation time can be expressed spatially in terms of the position of the target at movement initiation, and timing error in terms of where the center of the bat strikes the target (Fig. 4A). To assess whether delays to initiation and arrival were volitional, we ran three additional assays immediately following the experiment. In the first assay, participants watched a dummy Certain (counterbalanced Uncertain) trial and were asked to drag the target to where it would have been when they left the cave. For the second assay, the target appeared in the center of the screen and participants were asked to click where on the target they were aiming to hit. In a final assay, participants indicated where on the bat they were attempting to hit the target. This enabled us to center the reported position on the target relative to the center of the bat for analysis. The same procedure was then repeated for the Uncertain (counterbalanced Certain) condition.

**Figure 4.**
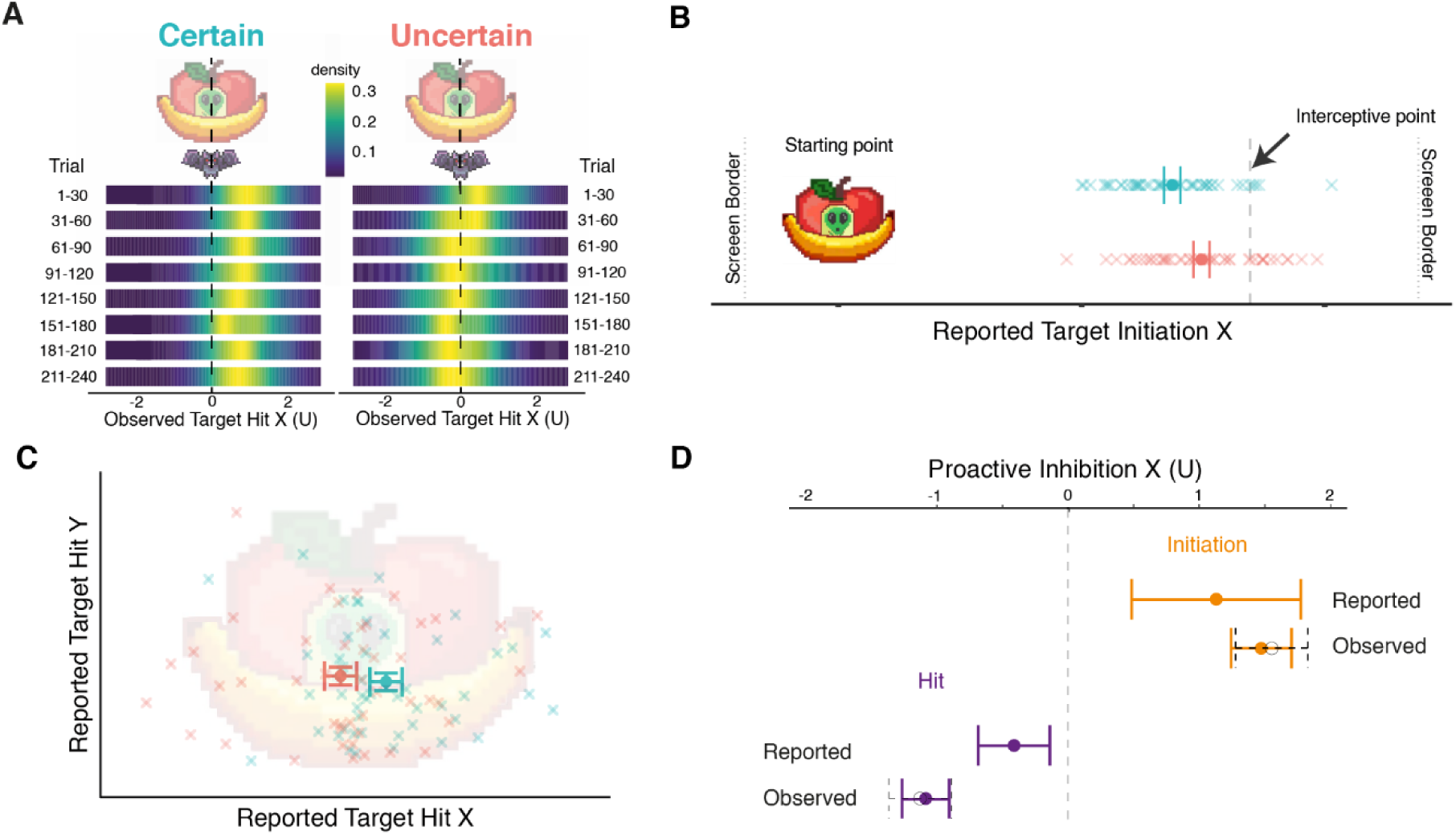
Spatial representation of observed timing errors and explicit reports of aiming strategies. A) Density plots of observed target hit x-coordinates relative to the center of the bat. In Certain conditions participants arrived at the front half of the target throughout the experiment. In Uncertain conditions, participants drifted towards the back half as the experiment progressed. B) Reported positions of target when participants initiated in Certain (blue) and Uncertain (red) conditions. Each cross represents a single participant, the dots (error bars) are the group means (95% within-subject adjusted CIs). Participants reported initiating when the target had travelled significantly further in the Uncertain condition. C) Reported position of where participants aimed to hit the target in each condition (normalized by reported bat position). Each cross represents a single participant, the dots (error bars) are the group means (95% within-subject adjusted CIs). Participants reported aiming significantly further to the back of the target in Uncertain conditions. D) Comparison between reported and observed differences in target initiation and arrival. For reported measures, dots (error bars) show mean spatial difference (95%CI) in reported initiation and hit between certainty conditions. For observed differences, coloured dots (error bars) show the mean difference (95%CI) from model estimates of initiation time (orange) and timing error (purple) at the end (i.e., the asymptote) of the experiment. Black dots (error bars) show the model-free mean differences (95%CI) observed in the final block. Positive (negative) values represent a target/hit position further to the right (left) in Uncertain conditions for initiation/arrival respectively. Participants reported delaying their arrival less than they delayed their initiation which corresponds to observed behaviour. Nevertheless, the differences observed were greater in magnitude than both reported measures.

Considering initiation, participants reported the target had travelled further before initiating in Uncertain relative to Certain conditions (*t*(48) = 3.53, *p* < .001, *d* = .50). Specifically, the target had travelled 3.34(± .48)U in the Uncertain condition compared to 2.22(± .54)U when Certain (Fig. 4B). Considering arrival, participants also reported aiming further to the rear of the target in Uncertain conditions(*t*(48) = 3.03, p = .004, *d* = -.43), with a mean bat-centred x-coordinate of -.14(±.23)U compared to .26(± 0.18)U when Certain (Fig. 4C). There was no difference in the reported y-position (*t*(48) = .66, p = .514).

Finally, to consider the extent to which explicit processes were responsible for the observed behaviour, we converted observed differences in initiation time and timing error at the end of the experiment into spatial units and compared them to the differences reported in the spatial assay (Fig. 4D). We observed the target having moved 1.47(± .23)U further prior to initiation in Uncertain conditions, whereas there was a reported mean difference of only 1.13(± .64)U. Furthermore, we observed participants arriving 1.08(± .18)U further to the rear of the target in Uncertain conditions, whereas the reported difference was only .41(± .27)U. Thus, the relative magnitudes of the explicitly reported strategies matched the observed behaviour, with participants reporting delaying their initiation more than their arrival, but the absolute magnitude of these effects was diminished compared to what was observed (Fig. 4D).

## DISCUSSION

Using a novel stop-signal task in which participants hit moving targets, we were able to probe the mechanisms underlying proactive inhibition. For the first time, we consider proactive changes to movement planning that are manifest downstream movement onset. We found delays to initiation were accompanied by shortened movement times and delayed arrival, suggesting both response suppression and response delay contributed. Moreover, participants reported strategies reflected this behaviour, suggesting explicit processes were responsible for proactive stopping.

Firstly, we validated our task, finding the assumptions made by the independent race model were met (Fig. 2A-D). In our task, a natural limit to initiation delays was enforced by having to hit a moving target. Thus it is likely we additionally avoided bias and faster preceding go-trials because they are mitigated by capping movement onset delays^28,30^. The average stop-signal reaction time was 197ms. This lies well within the measurement error of previous estimates in an interceptive timing context^9^ and aligns with the ∼200ms range observed in choice and simple response tasks. It should be noted it is considerably longer than the ∼150ms observed in anticipation timing tasks involving button presses^33^. This likely reflects the fact movements must be prepared further in advance when the movement time must be specified, with greater difficulties associated with inhibiting movements further in the preparatory stages^9^. Given the empirical assumptions were met and our estimates of SSRT were sensible, we are confident stop-signal uncertainty suitably constrained movements in our task.

Secondly, we considered proactive changes to the planning and execution of goal-directed movements. Under the uncertainty of whether a stop-signal could be presented, movement initiation was delayed. This has been previously observed in choice decision^16,34,35^ and anticipation timing^17,22,36^ contexts in which responses are recorded using simple button presses. Such proactive inhibition has been attributed to response suppression through activity in the circuitry responsible for reactive inhibition in anticipation of needing to stop^2^. An alternative explanation is that delays in our task simply reflected a “waiting strategy” in which participants delayed the facilitation of their response, withholding movement until certain a stop-signal would not be presented^23^. Resolving this issue is fundamental to our understanding of cognition. While there is a consensus neural inhibition is integral to the function and organisation of the brain, there is some doubt as to whether inhibition is implemented cognitively at all^20^.

The present study addresses this controversy by considering the movement time and timing error associated with delayed onset of the response. In the pre-programmed model of interception, the movement time is programmed immediately prior to the go signal triggering^9,24^. Therefore, if response suppression is implemented as an increase in the signal completion threshold^16^, movements will not be brief enough to preserve timing error because the movement duration cannot be updated once the plan has been triggered. Alternatively, any delays in facilitation can incorporate briefer movements because, by definition, they are triggered later.

In response to stop-signal uncertainty, we observed both briefer movements and delayed arrival, implying delays to initiation were achieved through a combination of a waiting strategy and response suppression. At the beginning of the experiment, these changes accounted for approximately equal proportions of the ∼100ms initiation delay (∼45ms briefer and ∼55ms later in arrival). By the end of the experiment, however, the ∼135ms initiation delay was offset by only ∼35ms briefer movements, with arrival delayed by ∼100ms. These data suggest waiting strategies are largely fixed, whereas suppression can be adapted between trials in response to feedback about performance, permitting even greater delays to initiation over many repetitions. In balance, our findings align with previous studies that favour response suppression as the mechanism driving delayed onset^21,35^, particularly as it pertains to increased delays over the course of the experiment^16^. Making the distinction between delays and suppression has important implications. Future research in clinical populations may more accurately characterise the cognitive deficiencies in neuropathologies such as Parkinson’s and ADHD, providing new insights into their development and treatment.

Of course, it is also possible that certain features of our task may have biased participants towards later arrival. For example, having been trained to move with a minimum movement time of 100ms, perhaps participants were conditioned not to shorten duration any further. We believe this is unlikely because at their quickest, participants were not close to approaching this limit (Fig. 3A), and there was no penalty or feedback in the event the limit was exceeded. Rather than not being able to shorten movement times, however, it could be simply that they did not need to. There was a relatively large timing window of approximately 400ms, thus it was not necessary to account entirely for the delay in initiation because it was also possible to delay the estimate of the moment of interception and still hit the target. While this has some merit, it does not explain why participants did not both aim later and shorten their movement time further, permitting greater delays to movement onset and further improvements to successful inhibition. On the other hand, the presence of response suppression does place a limit on the quickening of movement, because the movement duration must be long enough to hit the target in Certain conditions when initiation is earlier and the target is not directly overhead.

Finally, we considered whether these proactive changes were volitional. Our spatial assay allowed participants’ awareness of movement changes in movement planning to be quantified. In Uncertain conditions, participants reported initiating when the target was later in its trajectory (Fig. 4B) and aiming further to its rear (Fig. 4C). This supports the notion proactive inhibition is under volitional control^2,16,19^ because deliberate processes should be explicitly retrievable. However, we cannot entirely discount this was an explicit insight developed when questioned afterwards^37^. For example, the assay may have prompted retroactive retrieval of visual cues that coincided with performance, rather than the proactive strategy guiding it.

Still, an interesting feature was that both self-reported measures were smaller in magnitude than their observed counterparts (Fig. 4D). It is known that visual working memory for spatial estimates decays over time, with a reduction in the amplitude of spatial estimates when probed at increasing delays^38^. A reduction in magnitude of reported initiations and arrivals may simply reflect the fact participants were probed after the experiment concluded. This explanation is not entirely supported by the data, however, because there was no difference in reports between the counterbalanced orders of Certainty conditions, despite more time having elapsed once participants reported their second condition. It is therefore unlikely these reports simply reflected memory retrieval decay.

Assuming reports were of volitional strategies, it begs fundamental questions about the nature of the inhibitory process. For example, the underestimation of self-reported aim in reaching tasks has been accounted for by postulating an additional, implicit process that acts additively to produce the observed behaviour^32,39^. Analogously, proactive inhibition may be further reduced into implicit and explicit components. Specifically, there may be a categorical decision to implement a compensatory strategy whose details are implicitly tuned with experience (e.g., one may decide to delay initiation and adjust the magnitude of that delay based on feedback). While previous work has indirectly assessed the assumption that proactive inhibition is volitional and under conscious control^40^, our method is the first to quantify it, providing an avenue for disentangling the contributions of explicit and implicit processes underlying proactive inhibition. Indeed, given that we found response delay was relatively fixed, with an increase in response suppression over the course of the experiment, future research should consider the extent to which the division of implicit and explicit processes may follow these distinct dimensions.

In conclusion, these results demonstrate that in contexts where there is uncertainty over whether a movement needs to be withheld, there are proactive changes to the planning and execution of that movement beyond onset delay. In the context of timed interception, we found both shortened movement times and later arrivals, suggesting a combination of delayed facilitation and response suppression contributed. This has important implications for our understanding of inhibition because while the amount of inhibition will be correlated to onset delays in simple button press tasks, we have shown the degree to which response suppression and facilitation contribute is context dependent. Moreover, through quantifying the extent to which these changes were governed by explicit processes, we found participants were aware of both initiating and arriving later, but this did not account entirely for the difference observed. At bat, therefore, successful inhibition in response to an errant pitch may be augmented by a combination of delays and suppression to the planning and execution of the swing, improving the chances of detecting a need to stop. While this is underlain by explicitly retrievable processes, the extent to which this is the case, and how this corresponds to the underlying mechanisms, are important questions raised for future research.

## METHOD

### Participants

Sixty participants were recruited from Prolific, an online recruitment platform. All participants resided either in the UK or US, had English as their first language, and had a minimum Prolific approval rate 95%. Participants received £3.50 upon successful completion of the study, which took approximately 25 minutes to complete. Pay was contingent on completion and not performance. Three participants were omitted because they had more than 5 trials missing from their data. Additionally, as an online study, we set stringent screening requirements to ensure all participants included in the final sample completed the task as intended. Specifically, they had to hit more than 80% of targets in each condition, more than 80% of trials had to feature movement times less than 300ms, their probability of responding to a stop-signal must be between 40% and 60%, and the probability of omitting an appropriate go response must be less than 10%. Seven participants were excluded at this stage (4 for hit performance, 2 for elevated probability of stopping, and 1 for omitting too many go responses). Thus, the final sample for analysis was comprised of 50 participants (23 female, 4 left-handed, Mean_age_ = 30.56, SD_age_ = 9.90). The study was approved by the School of Psychology Ethics Committee at the University of Leeds (Reference: PSYC-116; Date: 27/10/2020).

### Experimental Task

Participants completed a browser-based game, *Fruit Bat Splat,* in which they had to hit targets moving across a screen with a bat that was controlled by the movement of their mouse or trackpad. The game was programmed in Unity (ver. 2019.4.4f1) by author JP and implemented in WebGL. Participants were not required to use standardized screen resolutions, sizes, or refresh rates. Accordingly, dimensions (in brackets) are given in Unity units (U). Unity units scale with the resolution of the screen. A reference resolution of 1920×1080 was used with screen dimensions of 17.75Ux10U with 1U corresponding to 108 pixels. Screens that were not native 16:9 were letterboxed to preserve the dimension of their longest side. Coordinates are defined with (0,0) at the center of the screen. Fullscreen was forced and the mouse pointer locked and hidden at the center of the screen throughout the experiment.

The position of the cursor was represented on screen by a pixel art fruit bat (1.75Ux1.5U), which had a fixed x-position of 5.5U but could move freely in the y-plane (Fig. 1). The bat’s wings were animated to improve participant engagement. Each flight cycle was comprised of four sprites: wings down, wings level, wings up, wings level. Each cycle lasted 20 frames (5 for each sprite). At the start of each trial, a start-zone represented by a pixel art cave (2.5Ux2.5U) appeared at the bottom right of the screen (5.5U, -4.5U). Participants were instructed to move the bat into the cave and click once. Subsequently, the bat snapped to the center of the start-zone and the flight animation was replaced with a static resting sprite, a bat hanging upside down from its feet inside the cave. On the same frame update, a target (3Ux2.25U) - depicted as a pixel art alien commandeering an Unidentified Fruity Object (UFO) – began to appear in the upper-left of the screen (−6.5U, 3.5U).

The appearance of the UFO was animated, so that it was fully visible after 350ms. After a short pause, randomly drawn from a uniform distribution (500-1500ms), the target began to move along the x-axis towards the vertical plane of the bat at 10U/s. The interceptive point, where the center of the target crossed the x-position of the bat, was in the upper-right of the screen (4.5U, 2.5U). The target reached the interceptive point after it had moved 11U along the x-axis, the bat reached the interceptive point after it had travelled 5U along the y-axis (Fig. 1). The bat’s flight animation would resume once it left the cave. If participants left the cave before the target began to move the target would disappear and the trial began again.

The study featured two certainty conditions: Certain and Uncertain (Fig. 1). On Certain trials, depicted by a task background of a mountain scene at night, participants were told it was safe for bats to leave the cave and to hit the target as it flew overhead (Certain Go*-*trials). On Uncertain trials, depicted by the same background mountain scene but at dawn, one third of trials presented a stop-signal before the target passed overhead. This was represented by the same scene instantly switching to daytime. Participants were told fruit bats burn easily in the sun and should try to withhold their movement in response to the stop-signal (stop-trials). But it was safe to leave the cave at dawn and to try to hit the target when a stop-signal was not presented (Uncertain go-trials). Uncertain trials were used to calculate our measure of reactive inhibition, Stop-signal Reaction Time (SSRT). Differences in *initiation time, movement time,* and *timing error* between Certain *go-*trials and Uncertain *go-*trials provide our three measures of proactive inhibition.

On go-trials, a successful hit could only occur if the upper edge of the bat contacted the lower edge of the target, indicated by a “HIT” graphic appearing in the center of the screen and accompanied by a splat sound effect. If the upper edge of the cursor moved past the lower edge of the target 1.5U before or after the center of the target had passed, the target stopped at that instant and a “MISS” graphic was displayed in the center of the screen. This prevented moving to the interception point early to “catch” the target.

On stop-trials, the stop-signal was first presented 320ms before the center of the target reached the interceptive point (i.e., a stop-signal delay of 780ms). This timing was derived from the mean stop-signal delay observed in a pilot study, ensuring the staircasing began near to the mean point at which movement is successfully inhibited 50% of the time. If the bat left the cave (i.e., displaced 0.75U), an “OUCH!” graphic appeared in the center of the screen, accompanied by a short animation of the bat being burnt in the sun and scorched sound effect. On the subsequent stop trial, the stop-signal delay was staircased 50ms shorter. If movement was successfully inhibited, a “SAFE!” graphic would appear in the center of the screen, accompanied by a cheerful chirping sound effect. On the subsequent stop trial, the stop-signal delay would be 50ms longer.

Feedback on all trials was displayed until participants clicked their mouse to advance to the next trial. For left-handed participants, the x-coordinates of the cursor and the target were reversed, with all sprites flipped horizontally.

### Procedure

Prior to the experiment, participants indicated whether they were using a mouse or trackpad to control the cursor. Then they calibrated their cursor to ensure movements were approximately equivalent between individuals. Using a sensitivity slider, participants reported when a 5U displacement of the bat corresponded to a “single flick” (mouse) or “single vertical stroke” (trackpad). This mapping was subsequently used throughout the experiment.

Participants were then given step-by-step instructions on completing the task (Supplement A). To ensure movements were rapid, participants were trained to move at a movement time between 100ms and 300ms and were unable to proceed to experimental trials until they had performed 5 successive movements within this movement time window. Participants were asked to move within this movement time window throughout the experiment, but this information was withheld so participants were not discouraged from delaying their responses and the effects of proactive inhibition could be observed.

Experimental trials were presented in 8 blocks of 30. Each block contained 15 Certain trials and 15 Uncertain trials. In Certain conditions, all 15 trials were go-trials. Of the Uncertain trials, 5 in each block were stop-trials and 10 were go-trials (Fig. 1D). The order of trials was randomized within each block. There were 240 trials in total. Participants were able to exit full screen, pause the game, and take a break at any time. Participants were unable to advance the experiment until they unpaused the game which once again forced full screen and locked the mouse pointer.

Following the main experimental blocks, we included two short tasks to investigate whether participants were explicitly aware of any differences in their movement characteristics between Certain and Uncertain conditions. Participants watched a simulated trial in which there was no bat to control, with either the Certain or Uncertain background. Before each trial, they were told to “*Pay attention to where the target is when you would leave the cave to hit it*” After the target travelled the full length of the screen it reappeared at its start point (−6.5U, 3.5U) with the instruction, “*Drag the UFO to the point you tried to leave the cave at [day/night]*”. Participants then dragged and dropped the target with their mouse to indicate where the target was when they initiated their movement. Next, the target and bat appeared in the center of the screen along with the instructions “*Click the point of the UFO you tried to hit at [day/night]*” and “*Click the point of the BAT you were using to hit the UFO at [day/night]*” They clicked where on the target they were aiming to hit and where on the bat they were using to hit, respectively. The above was subsequently repeated with the alternate certainty condition. The order was counterbalanced with half the participants reporting Certain first. The movement initiation report always came before the aiming reports in both conditions.

All task code can be found in an online repository at: https://github.com/jpickavance/response-inhibition-game-unity-project.

A demonstration version of the task can be found here: https://www.fruitbatsplat.com/demo

### Kinematic Analysis

All analyses were conducted after experimental data had been collected. Firstly, the position time series from each trial was resampled to a standard frequency of 100Hz. This is because Unity samples input at the same frequency as the monitor’s refresh rate which was beyond our control. 100Hz was chosen as the nearest round number between the most common framerates observed, 60Hz and 144Hz. We filtered the data using a zero-lag, 1^st^ order, low-pass Butterworth filter with 20Hz cutoff. The *initiation time* (IT) was calculated as the initial time at which the bat had moved 0.75U (i.e., the lower most edge of the bat had crossed the center of the start-zone). When considering proactive inhibition only, initiation times were converted to time to contact (TTC) initiation times by subtracting the observed initiation time from 1100ms. The resulting TTC initiation time thus expresses the time prior to the center of the target passing the interceptive point the initiation is made, with larger values earlier, and smaller values later in the target’s trajectory. The *movement time* (MT) was calculated as the time elapsed from the initiation time to the point at which the bat crossed the interceptive plane. We used cubic interpolation on all positions of the center of the target and the center of the bat, from initiation time until max movement amplitude, to estimate the precise moment the bat’s center intersected the interceptive plane (i.e., the y-position of the lowermost edge of the target). The *timing error* (TE) was calculated by dividing the displacement between the centers of the target and bat and dividing by the speed. Positive timing errors indicate the center of the bat arrived later than the center of the target, negative timing errors earlier.

### Stop-signal Reaction Times

Stop-signal Reaction Times (SSRTs) were estimated from Uncertain trials only. They were calculated for each participant using the integration method. Initiation times on go-trials were ranked and the *n*th trial was found where *n* is the total number of go-trials multiplied by the probability of responding. Individuals’ mean stop-signal delays (SSD) were subtracted from their go-trial initiation times on the *n*th trial to find their SSRT. This is suitable when using a staircasing procedure with at least 40 stop trials and their incidence is 33% or less^41^. Trials on which go responses were incorrectly omitted were assigned the maximum initiation time observed^28^. Trials were then ranked according to their initiation time. The *n*th IT was selected where *n* is the probability of responding given a stop-signal multiplied by the number of go responses observed. The SSRT was found by subtracting the mean stop-signal delay from the *n*th IT^41^. Actual values of the SSD were used to compute the mean (cf. programmed values) as the precise presentation of the stop-signal depended on the refresh rate of the display^28^. Summary statistics of the variables used to calculate SSRTs are displayed in *Table 1*.

### Proactive Inhibition

Differences in *initiation time*, *movement time,* and *timing error*, on go-trials between Certain and Uncertain conditions represent our measures of proactive inhibition. Additionally, when considering the effects of proactive inhibition, we wanted to ensure participants were timing their initiations to move with ballistic movements. Accordingly, trials were removed in which participants initiated before the target had crossed the halfway point, after the target had left the screen, or with a movement time of more than 300ms. In total, 621/9896 (∼6.3%) trials were removed at this stage.

Summary plots revealed each measure was characterized by a non-linear change that appeared to become asymptotic as trials progressed. Rather than take a single mean value for each participant, therefore, we modelled each dependent variable using a simple 3-parameter asymptote model:

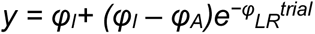

Where *φ_I_* is the y-intercept (i.e., value at beginning of experiment), *φ_A_* is the asymptote (i.e., approximate value at end of experiment), and *φ_LR_* is the learning rate. Prior to modelling, movement times were additionally transformed using a log transformation to normalize a positively skewed distribution.

To determine the significance of the effect of certainty at the start and end of the experiment, we began with a null model which computed the intercept on each parameter, with random intercepts for *φ_I_* and *φ_A_* for each participant. The effect of certainty on *φ_A_* (i.e., the asymptote) was determined by constructing a model which additionally included the effect of certainty on *φ_A_* with random slopes for each participant on *φ_A_*. A likelihood ratio test was then performed between the two models and the p-value taken as the level of significance of the effects of certainty on *φ_A_*^42^. We additionally considered a third model, including the effects of certainty on *φ_I_* (i.e., the intercept) and random slopes for each participant on *φ_I_*. A likelihood ratio test was performed between this and the best performing, most parsimonious model thus far (i.e., the asymptote model if significant, the null model if not). Finally, we allowed *φ_LR_* (i.e. the learning rate) to vary by certainty. Though we were primarily interested in estimating mean values at the start and end of the experiment (i.e., *φ_I_* and *φ_A_*), we wanted to ensure we had the most accurate estimates possible. Finally, this model was compared to the best performing model to point. Models were constructed using the *lme4* package in R studio (ver 3.6.0). Starting parameters were determined graphically. Optimisation was performed using the Broyden–Fletcher–Goldfarb–Shannon algorithm as implemented in the *optimx* package. The best performing model for all three measures included random slopes and intercepts on *φ_I_* and *φ_A_*, but no effect of certainty on *φ_LR_*, These comparisons can be found in Supplement B.

From the best model, individual fits for each participant were used to provide mean estimates for each dependent variable at the start and end of the experiment. We performed repeated-measures ANOVAs on these estimates to determine the significance of the effects of proactive control over the course of the experiment on each dependent variable. Shapiro-Wilk tests and qq-plots confirmed normally distributed data and variance for each of the contrasts. Prior to this, movement times were additionally transformed to their original scale by finding the exponent of the estimate. Individual model fits for each participant can be found in the supplementary materials. Finally, difference scores for IT, MT, and TE, were calculated per participant for each model, which represent our final measures of proactive control. The group means (95%CIs) are displayed for each measure at both the start and end of the experiment in *Fig. 3B*. Finally, to compare these effect sizes to explicit reports, we transformed estimates of IT and TE into target position at initiation and arrival respectively by multiplying by a factor of .011 (i.e., the distance the target travelled in Unity units (U) in 1ms; *Fig. 4D*). All data are reported as mean ±95%CI across all subjects.

### Model-free analysis

We also considered the mean of each measure for each participant over the first 30 (start) and final 30 (end) trials to sense check our models. The group means (95%CIs) are presented in *Fig. 3B* and *Fig. 4D* as black, hollow dots (dotted lines). Differences between model and model-free estimates were negligible, though the modelled estimates provided more precision with smaller CIs. All data are reported as mean ±95%CI across all subjects.

### Reported Aim

We had two explicit measures of proactive control in each condition. *Reported target position* is how far the target had travelled when participants reported they tried to initiate their movement in each condition. *Reported target hit* is where on the target participants reported aiming to hit the target in each condition. The latter was centred by subtracting away the point of the bat they reported using to hit the target. One participant was omitted from this stage of analysis (failed to provide any data), with 49 participants considered in total. 2 x 2 mixed ANOVAs confirmed the certainty order in which participants reported their aiming strategy had no effect on either *reported target position* (F(1, 47) = . 026, p = .873) or *reported target hit* (F(1,47) = .738, p = .394), nor did it interact with certainty (F(1, 47) = . 389, p =.535; F(1,47) = .537, p =.467). Accordingly, data was combined, and paired sample t-tests were run on both dependent variables between the two certainty conditions. This was to determine whether participants were actively aware of adjusting their initiation or arrival in response to uncertainty. Difference scores between Certainty conditions for each participant were also calculated which are displayed at the group level in *Fig. 4D*. All data are reported as mean ±95%CI across all subjects.

## ACKNOWLEDGMENTS

We would like to thank Ian Greenhouse for his critical reflections on a draft version of this manuscript.

## AUTHOR CONTRIBUTIONS

J.P.P. and J.R.M. conceived and designed the research; J.P.P. programmed the task; J.P.P. built online systems for task hosting and database; J.P.P. ran the statistical analysis; J.P.P. and J.R.M. interpreted the results; J.P.P. prepared the figures; J.P.P. drafted manuscript; J.P.P., F.M., and J.R.M. edited and revised manuscript; J.P.P., M.M-W., F.M., and J.R.M approved final version of manuscript.

## Notes

### Competing Interest Statement

The authors have declared no competing interest.

