## Supplemental Materials for "Proactive inhibition of goal-directed movements involves explicit changes to movement planning"

### SUPPLEMENTARY MATERIALS

#### Supplement A: Task Instructions

*Instructions are given in italics*

[requirement to advance is given in parentheses – if requirement is not met, trial repeats]

**Main task training**

1. *Use the mouse to move the cursor into the cave (blue/night background displayed - Certain Go-trial)*
   [bat over cave]

2. *The cave turns green when the bat enters. CLICK the mouse to rest*
   [click mouse]

3. *Keep still and wait for the target to fly closer*
   [movement withheld]

4. *Wait for the fruit to fly over the cave. Try to hit it with the bat!*
   [hit the target with the bat]

5. *The fruit won't always be this slow*
   [click to advance feedback and cue a faster moving target]

6. *Try to hit the fruit again (Certain Go-trial)*
   [hit the target with the bat]

7. *Fruit bats get tired: flying too fast OR flying too slow. Use the flight timer to help you avoid getting tired (Certain Go-trial)*
   [hit the target with the bat and a movement time between 100-300ms]

8. **Movement time test.** *Hit the fruit. Move with a good flight time. Goal: 5 in a row (Certain Go-trial)*
   [hit the target with a movement time between 100-300ms 5 times consecutively. Score is reset to zero after a movement quicker than 100ms or slower than 300ms]

9. **Movement time training passed.** *Congratulations, you have completed the first round of training! Your excellent performance means you will no longer have the flight timer to help you. The flight time will still be recorded so try to fly with a good flight time in all future rounds.*
   [click mouse]

10. *Fruit bats easily sunburn. Stay inside the cave during the day (white/day background displayed)*
    [movement withheld]

11. *Try to hit the fruit with a good flight time at night. Stay inside the cave at day*
    [hit target on night trials, withhold movement on day trials – 3 of each in total]

12. *Fruit bats can still collect fruit at sunrise. Hit the fruit with a good flight time* (background changes to red/dawn *Uncertain Go-trial)*
    [hit the target with the bat]

13. *At sunrise, it can turn to day. Stay inside the cave if it turns to day. (Uncertain Stop-trial with SSD of 580ms)*
    [movement withheld]

14. **Stop signal test** *Hit the fruit with a good flight time at night. Hit the fruit with a good flight time at sunrise. Stay inside when it turns to day.*
    [Randomly interleaved trials:
    3 x Certain Go trials: hit the target with the bat
    4 x Uncertain go trials: hit the target with the bat
    4 x Uncertain Stop trials: withhold movement]

15. **Stop signal training passed.** *You have completed training*
    *The rules are the same for the full game: Night- hit the fruit with a good flight time. Sunrise- hit the fruit with a good flight time. Day- stay inside the cave. There will be 240 trials in total. You will be graded on your performance at the end. Good luck!*

**Warning messages during task**

**Movement > 300ms:** *You moved too slow. Try not to leave the cave until you can hit the fruit with one smooth movement*

**Movement withheld on go trial:** *You did not move. Please try to hit the fruit at night and sunrise*.

**Self-reporting initation and aim instructions**

1. *You will now see the UFO fly overhead. Pay attention to where the target is when you would leave the cave to hit it.*

[click ready]

2. *Drag the UFO to the point you tried to leave the cave at (day/night)*
   [drag and drop UFO to point on screen and click submit]

3. *Click the point of the UFO you tried to hit at (day/night)*
   [click on the UFO and then submit]

4. *Click the point of the BAT you were using to hit the UFO at (day/night)*
   [click on the bat and then submit]

#### Supplement B: Modelling

##### Method

Summary plots revealed each measure was characterized by a non-linear change that appeared to become asymptotic as trials progressed. Rather than take a single mean value for each participant, therefore, we modelled each dependent variable using a simple 3-parameter asymptote model:

$$y = \varphi_I + (\varphi_I - \varphi_A)e^{-\varphi_{LR}^{trial}}$$

Where $\varphi_I$ is the y-intercept (i.e., value at beginning of experiment), $\varphi_A$ is the asymptote (i.e., approximate value at end of experiment), and $\varphi_{LR}$ is the learning rate. Prior to modelling, movement times were additionally transformed using a log transformation to normalize a positively skewed distribution.

To determine the significance of the effect of certainty at the start and end of the experiment, we began with a null model which computed the intercept on each parameter, with random intercepts for $\varphi_I$ and $\varphi_A$ for each participant. The effect of certainty on $\varphi_A$ (i.e., the asymptote) was determined by constructing a model which additionally included the effect of certainty on $\varphi_A$ with random slopes for each participant on $\varphi_A$. A likelihood ratio test was then performed between the two models and the p-value taken as the level of significance of the effects of certainty $\varphi_2$ (Winter., 2013). We additionally considered a third model, including the effects of certainty on $\varphi_I$ (i.e., the intercept) and random slopes for each participant on $\varphi_I$. A likelihood ratio test was performed between this and the best performing, most parsimonious model thus far (i.e., the asymptote model if significant, the null model if not). Finally, we allowed $\varphi_{LR}$ (i.e. the learning rate) to vary by certainty. Though we were primarily interested in estimating mean values at the start and end of the experiment (i.e., $\varphi_I$ and $\varphi_A$), we wanted to ensure we had the most accurate estimates possible. Finally, this model was compared to the best performing model to point. Models were constructed using the *lme4* package in R studio (ver 3.6.0). Starting parameters were determined graphically. Optimisation was performed using the Broyden–Fletcher–Goldfarb–Shannon algorithm as implemented in the *optimx* package.

**Results**

The initiation time asymptote was different across conditions of trial certainty($\chi^2(4) = 3973.8$, $p < .001$), as was the parameter for initiation time at the beginning of the task ($\chi^2(5) = 227.1$, $p < .001$). The learning rate parameter was not significant ($\chi^2(1) = .035$, $p = .853$). Therefore, estimates for individuals' initiation times at the beginning and end of the experiment were computed using the model with fixed and random effects of certainty on $\varphi_I$ and $\varphi_A$.

Our fitted parameters for movement time were affected by the trial certainty ($\chi^2(4) = 1444.7$, $p < .001$) and this also affected our parameter for movement times at the start of the task($\chi^2$ (5) = 260.1, $p < .001$) but not the rate of change throughout the task ($\chi^2(1) = 1.51$, $p = .219$). We therefore used estimates of individuals' movement times at the beginning and end of the experiment using the model with fixed and random effects of certainty on $\varphi_1$ and $\varphi_2$ (Fig. 3).

The effect of certainty on the asymptote of timing error was significant ($\chi^2(4) = 2753.7$, $p < .001$). In addition, the effects of certainty on the intercept were also significant ($\chi^2(5) = 54.17$, $p < .001$). Moreover, the effects on learning rate did slightly improve the model ($\chi^2(1) = 4.06$, $p = .0439$). Thus, the effects of proactive inhibition were examined using estimates of individuals' timing errors at the beginning and end of the experiment using the model with fixed and random effects of certainty on $\varphi_I$, $\varphi_A$, and $\varphi_{LR}$ (Fig. 3).

##### Individual fits

###### *Initiation Time*

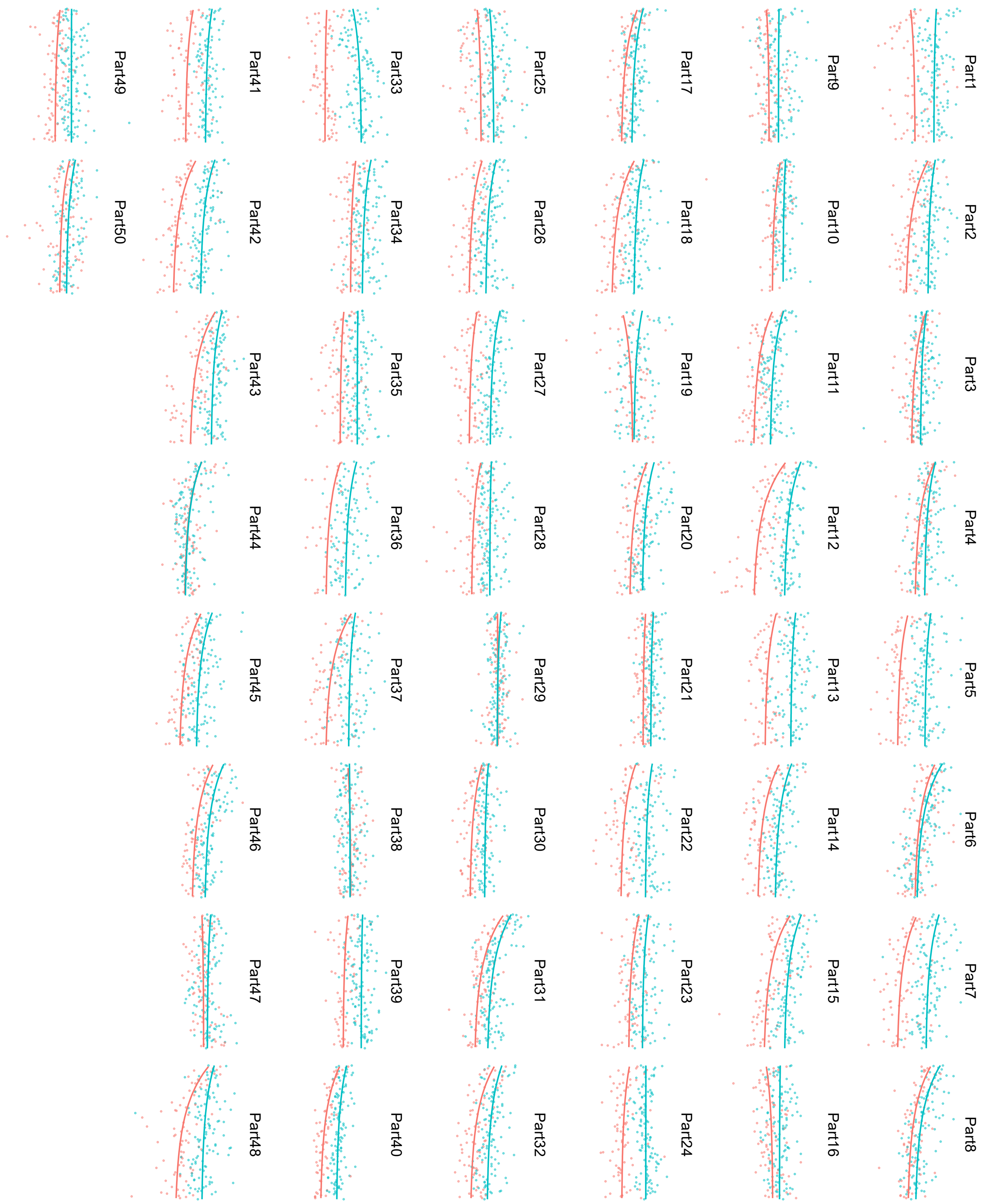

*Movement Time*

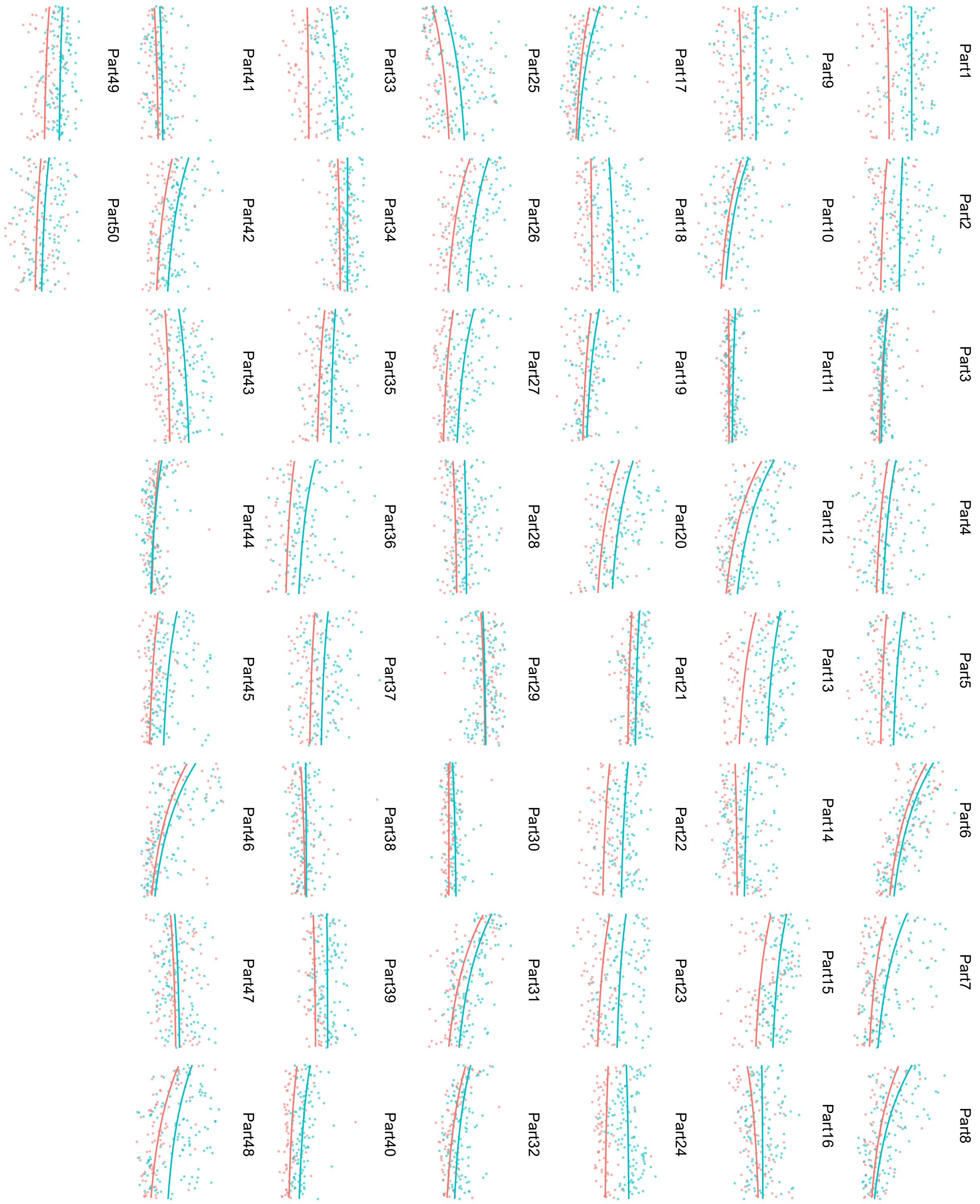

*Timing Error*

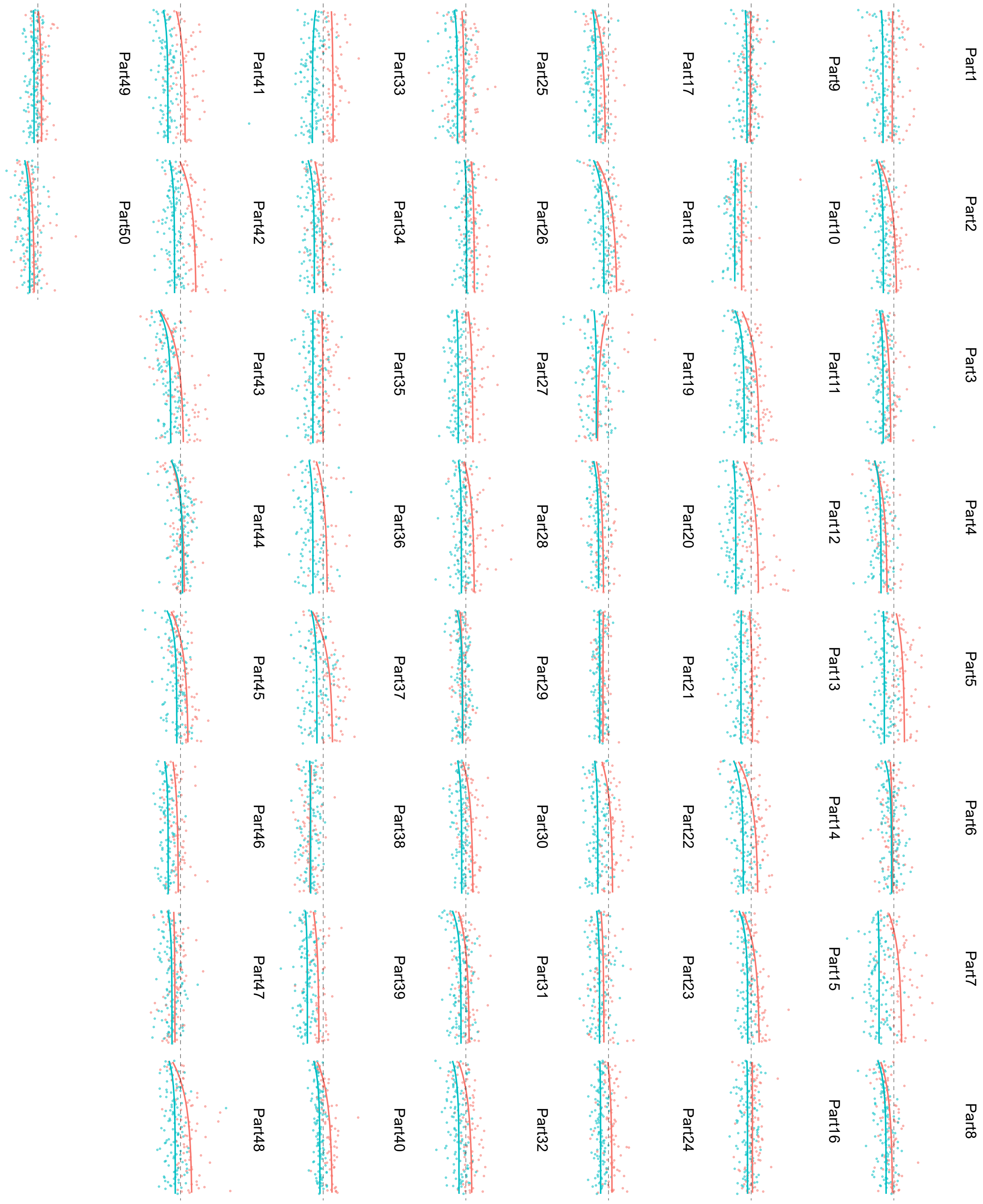

**Supplement C: Context Independence Violations**

Recent work has shown that go-trial initiation times can be quicker than stop-trial initiation times when they immediately precede them (Brisset et al., 2021). To see whether this was a problem in our task, we grouped individuals' stop-trials by (centered) stop signal delay. Considering only the initiation times of the immediately preceding go-trial on stop-trials there was an incorrect response, we found mean initiation times were quicker on stop-trials than on immediately preceding go-trials for all SSDs with more than 10 pairs of responses (Fig. A)

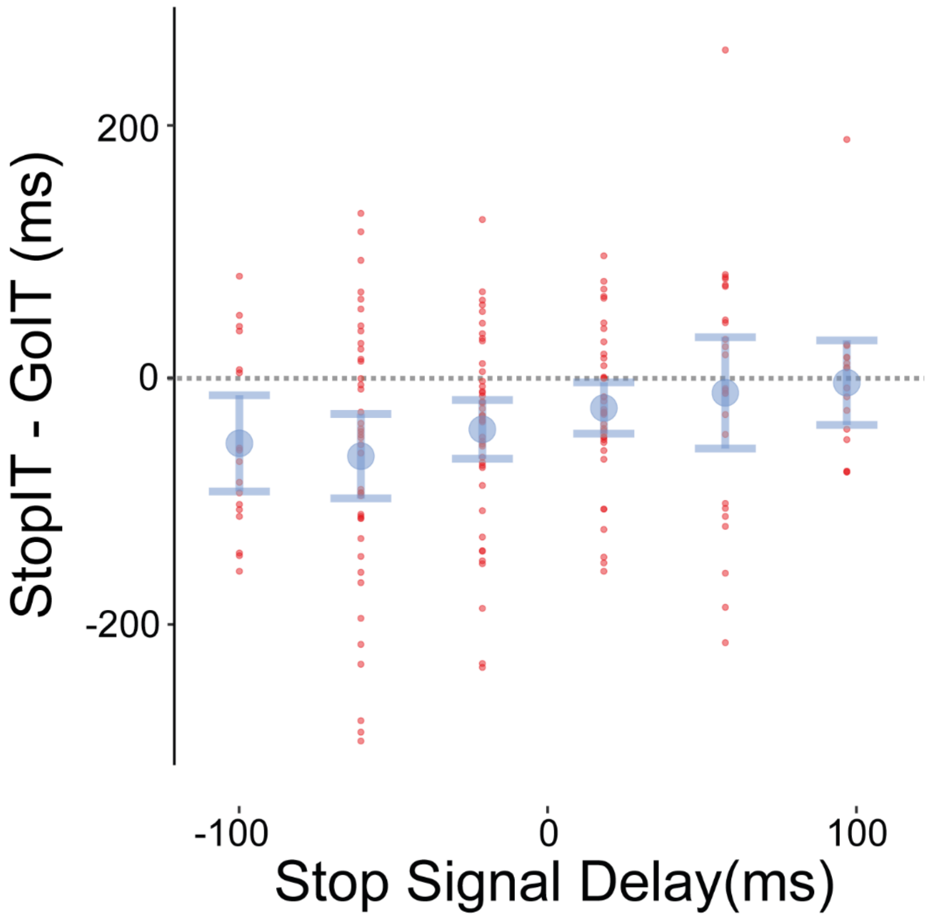


*Figure A* . Comparing the difference between stop-trial initiation times with go-trial initiations that immediately precede them. Red dots are participants average differences grouped by stop signal delay. Blue dots (error bars) represent the group mean difference(95% ci) at each stop signal delay. *For the sake of direct comparison, stop signal delays have been cantered for each participant by subtracting their mean SSD. Mean stop-trial initiation times are quicker, or no longer than mean go-trial initiation times at all levels of stop signal delay.*
